# A susceptibility locus on chromosome 13 profoundly impacts the stability of genomic imprinting in mouse pluripotent stem cells

**DOI:** 10.1101/2020.01.25.915413

**Authors:** Emily Swanzey, Thomas F. McNamara, Effie Apostolou, Mamta Tahiliani, Matthias Stadtfeld

## Abstract

Cultured pluripotent cells accumulate detrimental epigenetic alterations, including DNA methylation changes at imprinted genes known as loss-of-imprinting (LOI). Despite the substantial biomedical relevance of this phenomenon, the molecular cause of this epigenetic instability in pluripotent cells remains unknown. While the occurrence of LOI is generally considered a stochastic phenomenon, here we document a strong genetic determinant that segregates mouse pluripotent cells into epigenetically stable and unstable cell lines. Unstable lines exhibit hypermethylation at *Dlk1-Dio3* and select other imprinted loci, which is associated with impaired developmental potential. Stimulation of demethylases by ascorbic acid prevents LOI and can preserve developmental potential. Susceptibility to LOI greatly differs between commonly used mouse strains, which we utilize to map a causal region on chromosome 13 with Quantitative Trait Locus (QTL) analysis. Our observations identify a strong genetic determinant of locus-specific epigenetic abnormalities in pluripotent cells and provide a non-invasive way to suppress them. This highlights the importance of considering genetics in conjunction with culture conditions for assuring the quality of pluripotent cells for biomedical applications.

## Introduction

Genome-wide remodeling of DNA and histone marks plays a crucial role in establishing pluripotency in the pre-implantation mammalian embryo (Cantone and Fisher, 2013). A permissive chromatin structure resembling the early embryo is also a characteristic of embryonic stem cells (ESCs) and induced pluripotent stem cells (iPSCs) (Tee and Reinberg, 2014). While this unique chromatin state likely contributes to the broad developmental potential of these cells, ESCs and iPSCs are also susceptible to accumulation of epigenetic alterations during culture (Bar and Benvenisty, 2019). Some of these alterations, such as changes in genome-wide DNA methylation levels, remain developmentally controlled and reversible (Pastor et al., 2016), but prolonged culture can result in irreversible erosion of vital epigenetic control mechanisms at specific gene loci and concomitant loss of developmental potential (Choi et al., 2017; Yagi et al., 2017).

In particular, irreversible abnormalities have been reported at imprinted genes in cultured pluripotent cells. Imprinted gene loci are characterized by differential epigenetic modifications, often DNA methylation, that are established in sperm and oocyte at imprinting control regions (ICRs) and determine allele-specific expression patterns of nearby genes after fertilization (Duffie and Bourc’his, 2013; Sanli and Feil, 2015). Dysregulation of imprinted gene loci by aberrant loss or gain of DNA methylation – referred to as loss-of-imprinting (LOI) – is associated with severe developmental defects in mice derived from pluripotent cells (Dean et al., 1998; Stadtfeld et al., 2010; Stelzer et al., 2015). Importantly, once lost, the allele-specific expression of imprinted genes cannot be readily restored (Dirks et al., 2019; Pastor et al., 2016), making LOI a significant hurdle for the biomedical use of ESCs and iPSCs.

While numerous studies have characterized the degree of LOI in pluripotent cells, little is known about the underlying molecular mechanism. Based on line-to-line variability, LOI has been proposed to be a stochastic event triggered by a general epigenetic instability of pluripotent cells (Humpherys et al., 2001; Johannesson et al., 2014). However, specific imprinted gene clusters such as *DLK1-DIO3* and *IGF2-H19* in human ESCs, have been shown to be particularly susceptible to LOI (Kim et al., 2007; Rugg-Gunn et al., 2007), suggesting the presence of unknown molecular determinants. In the mouse, the *Dlk1-Dio3* ICR, known as the intergenic differentially methylated region (IG-DMR), can become methylated on the maternal allele during iPSC derivation, resulting in LOI (Carey et al., 2011; Stadtfeld et al., 2010; Stadtfeld et al., 2012). The imprinting status at this cluster also correlates with developmental potential in mouse and human pluripotent cells and may serve as an indicator of pluripotent cell quality (Liu et al., 2010; Mo et al., 2015).

In this study, we have used LOI at *Dlk1-Dio3* as a paradigm to investigate the mechanisms of epigenetic instability of pluripotent cells. For this, we took advantage of an allele-specific fluorescent reporter system that allows assessment of *Dlk1* imprinting without the need for more cumbersome single nucleotide polymorphism (SNP) sequencing (Swanzey and Stadtfeld, 2016). We confirm a high degree of variability of imprint stability between different ESC lines, but show that this variability is not stochastic in nature. Rather, it is caused by genetic determinants that establish a dynamic epigenetic state prone to locus-specific DNA hypermethylation in susceptible cell lines. This susceptibility can be suppressed by culture conditions that favor activation of enzymes involved in histone and DNA demethylation. Widely used mouse strains such as 129SvImJ (129) and C57BL/6J (B6J) differ in their susceptibility to LOI and we map a genomic region that determines this susceptibility. Our observations reveal the importance of genetic factors in conjunction with environmental conditions to control the epigenetic stability of developmental regulators in pluripotent cells.

## Results

### Variable stability of *Dlk1-Dio3* imprinting is observed in early passage ESC lines

We have previously described a transgenic system that allows assessment of *Dlk1-Dio3* imprinting based on allele-specific fluorescent reporter genes inserted into *Dlk1* (Swanzey and Stadtfeld, 2016). In brief, if LOI has occurred and the IG-DMR is hypermethylated, *Dlk1* reporter fluorescence from both parental alleles is visible (Fig. 1A). In contrast, retention of parent-of-origin methylation, known as maintenance of imprinting (MOI) results in detection of only the fluorescence from the paternally-inherited allele (Fig. 1A). To study the stability of *Dlk1-Dio3* imprinting status in cultured pluripotent cells, we established a cohort of ESC lines carrying this dual-reporter system. Flow cytometry evaluation revealed a large degree of variation in allele-specific *Dlk1* expression between lines as early as passage 6 (p6) (Fig. 1B, C). Mean CpG methylation at the IG-DMR was consistent with the percentage of cells expressing bi-allelic *Dlk1* (Fig. 1D), underscoring the reliability of the reporter system in measuring DNA methylation levels (Swanzey and Stadtfeld, 2016). Functional assessment by tetraploid (4N) blastocyst injection showed that ESCs with a low degree of MOI (ES6) exhibited strongly reduced developmental potential compared to ESCs with high degree of MOI (ES10) (Fig. 1E). Together, these observations demonstrate variable stability of *Dlk1-Dio3* imprinting and associated differences in developmental potential of ESCs derived under identical conditions.

**Fig. 1.**
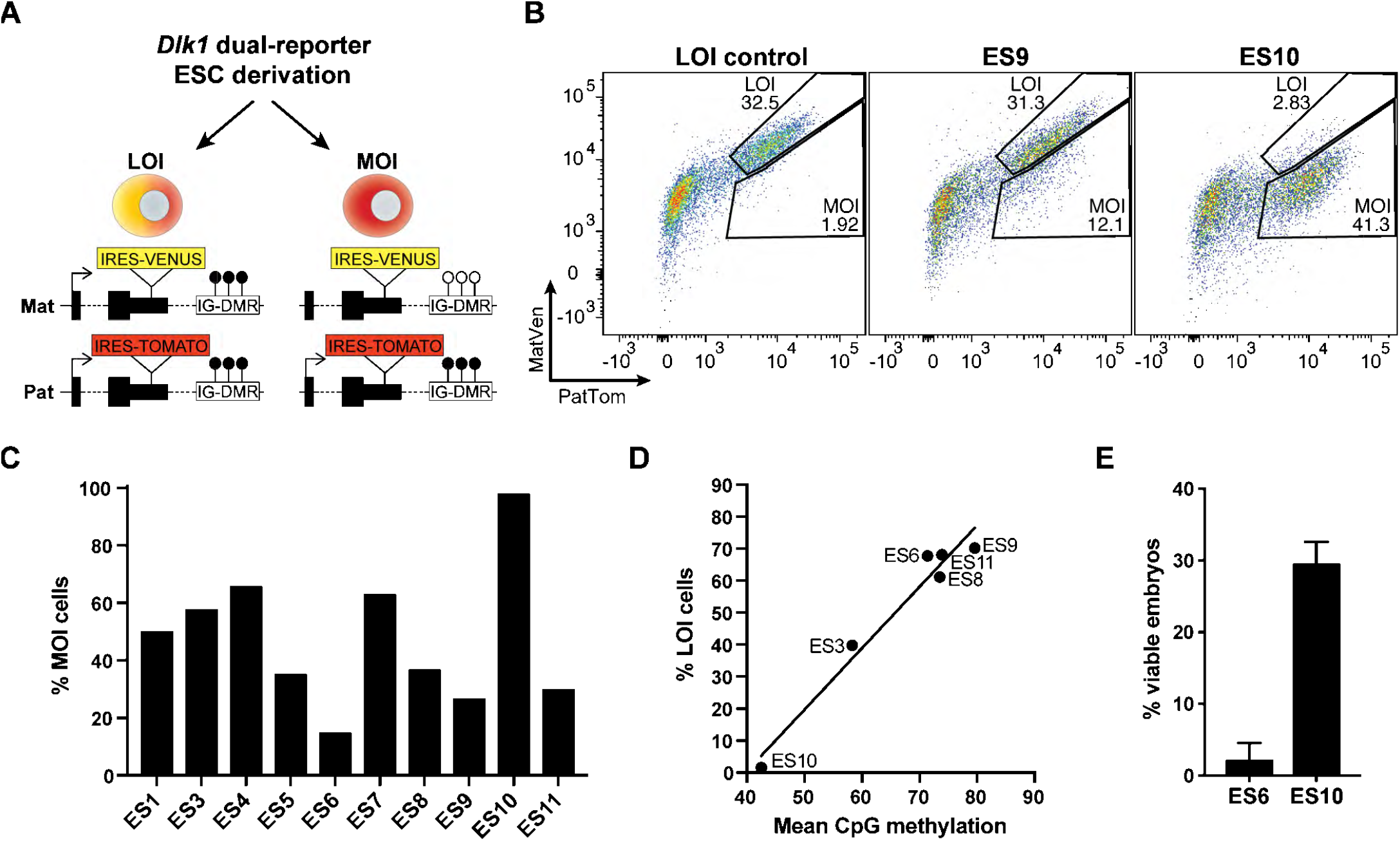
Variable stability of *Dlk1-Dio3* imprinting is observed in early passage ESC lines. (A) Schematic of the *Dlk1* reporter system with IRES-tdTomato (Tomato) on the paternal (pat) and IRES-Venus on the maternal (mat) allele and expected fluorescence and methylation in LOI and MOI cells. Lollipops indicate methylation status of the IG-DMR and an arrow represents active transcription. (B) Flow cytometric analysis of ESC lines ES9 and ES10 at p6, along with an iPSC line as LOI control. MOI and LOI gates are indicated. Cells were exposed to retinoic acid to trigger differentiation and *Dlk1* expression in a reliably measurable fraction of cells. (C) Percentage of *Dlk1*-expressing cells that have MOI in ten reporter ESC lines. (D) Percentage of *Dlk1*-expressing cells that display LOI reporter fluorescence (y-axis), compared to mean CpG methylation at the IG-DMR (x-axis) in six reporter ESC lines. (E). Percentage of transferred blastocysts that resulted in viable E15.5 embryos after 4N complementation with p6 reporter ESCs with opposite imprint phenotypes (ES6 and ES10). A total of 44 (ES6) and 64 (ES10) blastocysts were transferred. Lines were karyotyped at the time of 4N injection and were considered normal, with 70% (ES6) and 73% (ES10) euploid cells (data not shown). Error bars represent standard error.

### Increase in LOI in unstable ESC lines is due to epigenetic instability

To better understand the dynamics of the observed imprint loss, we followed reporter ESC lines that were derived and cultured simultaneously from p2 to p16. Surprisingly, while the degree of MOI rapidly dropped in the vast majority of lines, rare lines (5 of 38 evaluated) retained almost exclusively cells with MOI (Fig. 2A). We will refer to lines exhibiting such phenotypes as *unstable* and *stable*, respectively (Fig. S1A). Neither differences in cell growth nor sex could account for the differences in imprint stability and we observed indistinguishable pluripotent cell colony morphology between stable and unstable lines (Fig. S1B,C and data not shown).

**Fig. 2.**
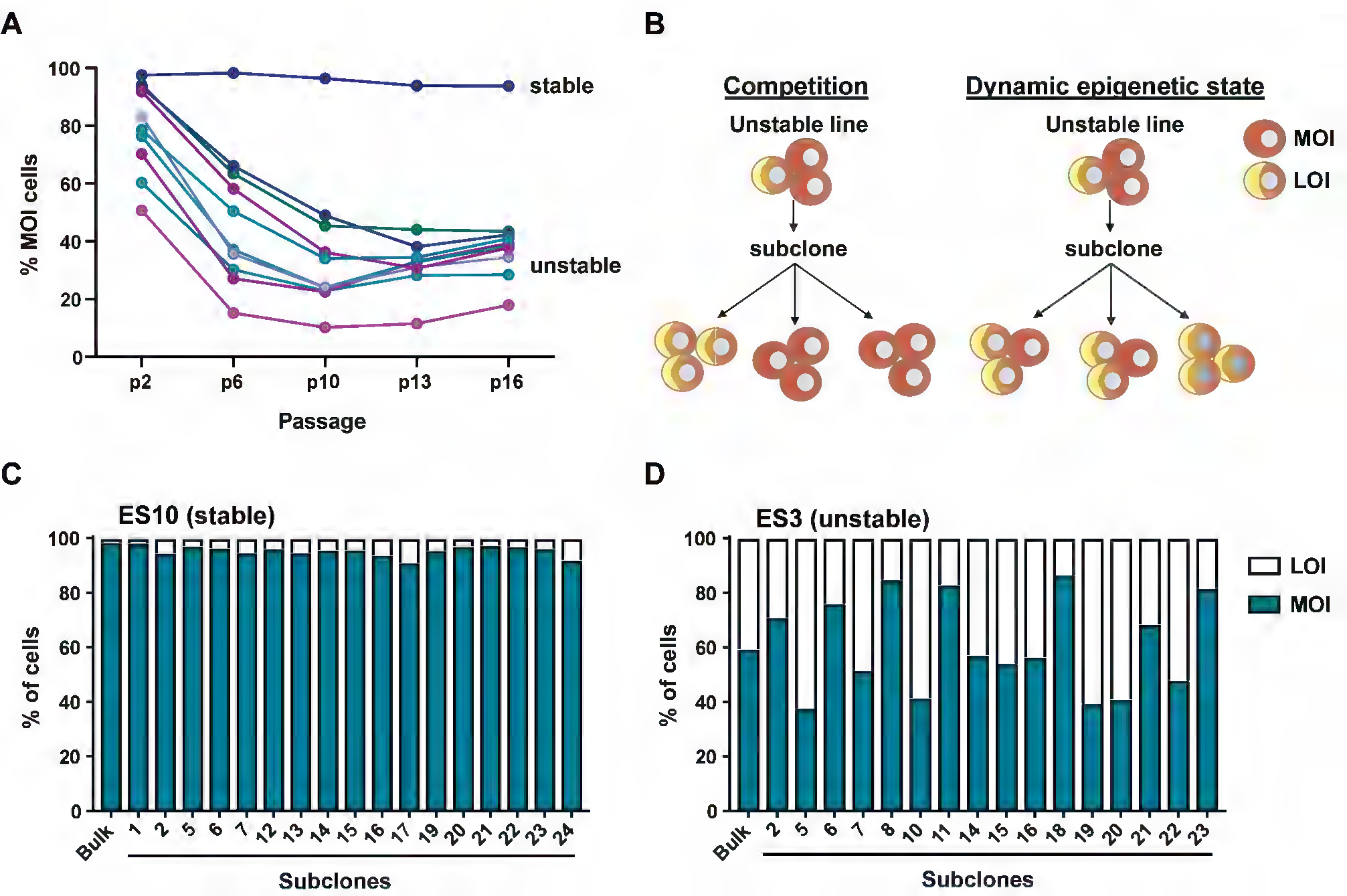
Increase in LOI in unstable ESC lines is due to epigenetic instability. (A) Percentage of *Dlk1*-expressing cells with MOI upon differentiating ten reporter ESC lines at the indicated passages (p2 to p16). Each colored line represents one cell line. (B) Schematic of the expected outcomes of subcloning an unstable line. In case of competition, subclones should have distinct expression patterns of either MOI or LOI. In case of a dynamic epigenetic state, subclones would be expected to resemble the heterogeneous expression pattern of their parental unstable line. (C, D) Single-cell subclones of one stable and one unstable line were analyzed by flow cytometry. Bar graphs show the quantification of the percentage of *Dlk1*-expressing cells that have LOI or MOI reporter fluorescence, compared to the bulk parental lines.

We reasoned that the gradual increase in cells expressing *Dlk1* in a bi-allelic manner in unstable lines could be the result of competition between MOI and LOI cells in a mixed culture, with LOI cells becoming predominant over time. Alternatively, our observations could indicate a dynamic epigenetic state that differs between stable and unstable lines (Fig. 2B). To distinguish between these possibilities, we generated subclones from two unstable lines and one stable line by single-cell sorting at early passage. Flow cytometry showed that subclones resembled the *Dlk1* expression pattern of their respective parental lines (Fig. 2C,D). This is inconsistent with the expectations of the competition model (Fig. 2B). Instead, our observations support the existence of a dynamic epigenetic state that favors gradual acquisition of DNA hypermethylation at the IG-DMR in unstable ESCs (Fig. 2B). We also noticed rare unstable lines that exhibited not only a high degree of biallelic *Dlk1* expression but also a population of cells with maternal-only expression (Fig. S1D), which was recapitulated upon subcloning (Fig. S1E,F). Together with the variable MOI observed at early passage (Fig. 1C) and the different kinetics of imprinting loss (Fig. 2A), this suggests a gradient of phenotypic severity in reporter ESC lines.

### Ascorbic acid attenuates imprint loss in unstable lines and prevents loss of developmental potential

Ascorbic acid (AA) is a known activator of histone and DNA demethylases (Cimmino et al., 2018) and a potent modulator of DNA methylation in ESCs (Blaschke et al., 2013). We therefore decided to test the effect of AA supplementation to the culture media of unstable reporter lines at early passage (see outline in Fig. 3A). Assessment by flow cytometry showed almost complete retention of MOI in unstable lines when cultured in the presence of AA (Fig. 3B). AA withdrawal at an intermediate passage (Fig. S2A) resulted in a pronounced decrease of MOI, suggesting that AA did not permanently correct the underlying molecular cause (Fig. S2B).

**Fig. 3.**
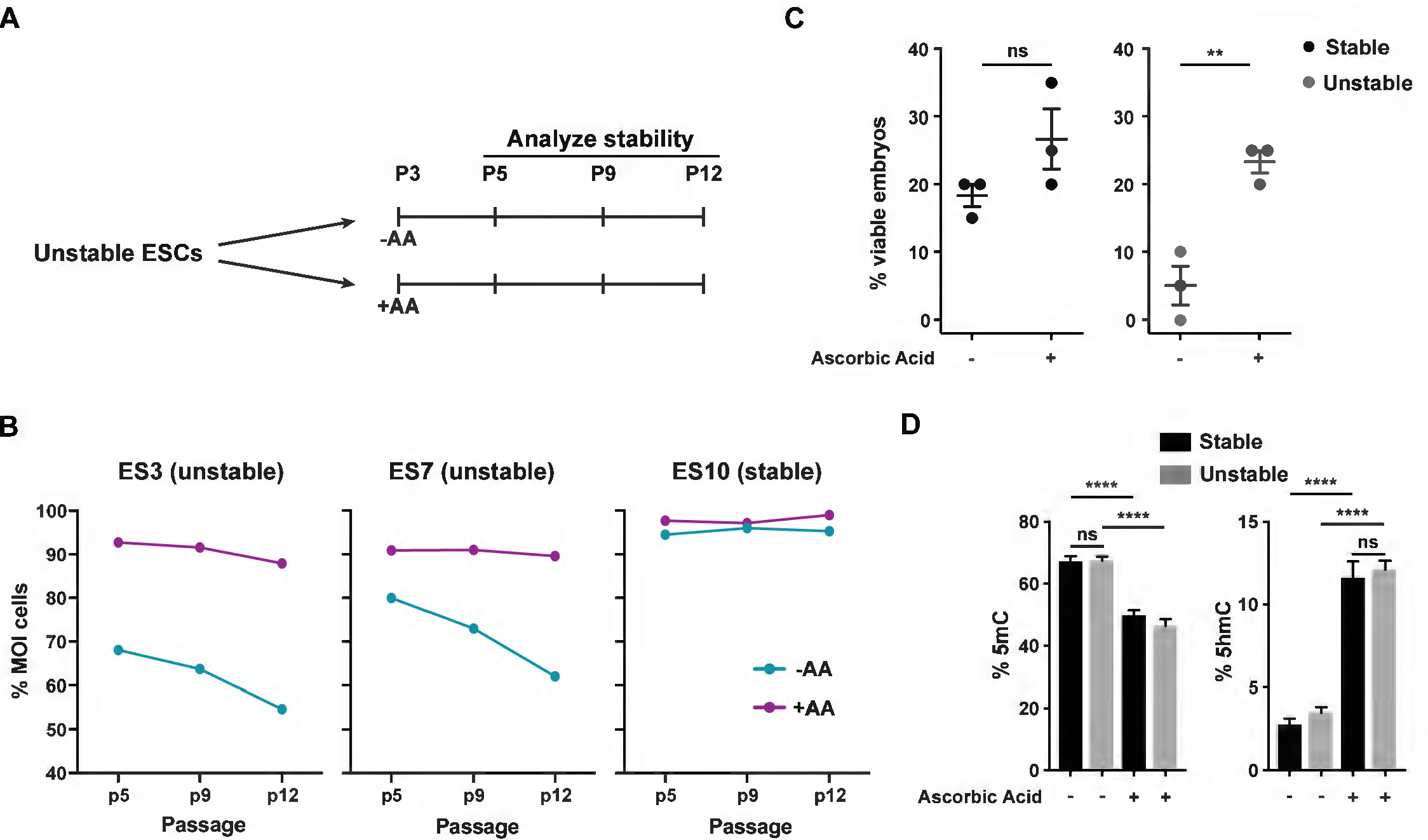
Ascorbic acid attenuates imprint loss in unstable lines and prevents loss of developmental potential. (A) Schematic of experimental approach. Unstable ESC lines cultured in standard conditions were split at early passage and either continued in the same conditions (−AA) or treated with AA (+AA). Imprint stability was assessed at indicated passages. (B) Percentage of *Dlk1*-expressing cells with MOI in two unstable (ES3 and ES7) lines and one stable (ES10) line in presence or absence of AA at the indicated passages. (C) Percentage of viable embryos at E14.5 after 4N complementation with stable (ES10) and unstable (ES3) lines cultured in presence or absence of AA from p2 to the time of injection (p7). Each dot represents one recipient female into which 20 blastocysts were transferred (with a total of 3 recipients per cell line). Bars represent mean and standard error. (**) p<0.01, (ns) not significant with a two-tailed students t-test. (D) Quantification of the percentage of 5mC and 5hmC as measured by TLC in stable and unstable ESCs (3 biological replicates per condition). Bars represent standard error. (****) indicates p<0.0001, (ns) not significant with a one-way ANOVA and Turkey’s multiple comparison test.

Importantly, we observed a significant increase in embryo viability after 4N blastocyst injections with unstable lines that were grown in presence of AA, with the overall efficiency of embryo recovery comparable to stable ESCs (Fig. 3C). Fluorescence microscopy revealed LOI in two of the three embryos that developed from unstable ESCs cultured in absence of AA, contrasting with embryos from stable ESCs and unstable cells treated with AA (Fig. S2C,D). This abnormality would have likely hindered further development of these embryos (Stadtfeld et al., 2010; Stelzer et al., 2015). Together, these observations indicate that AA can preserve the developmental potential of epigenetically unstable pluripotent cells by efficiently suppressing LOI.

Quantification of 5-methylcytosine (5mC) levels by thin layer chromatography (TLC) showed no significant difference between stable and unstable lines, suggesting that global methylation was not dysregulated in unstable cells (Fig. 3D and Fig. S2E). Given that AA is a cofactor of ten-eleven translocation (TET) proteins in the oxidation of 5mC into 5-hydroxymethylcytosine (5hmC) (Monfort and Wutz, 2013), a decrease in 5mC and an increase in 5hmC would be expected in the presence of AA if TET proteins are functioning normally. Indeed, stable and unstable lines displayed a similar increase in 5hmC and a concomitant drop in 5mC levels in presence of AA (Fig. 3D and Fig. S2E). Taken together, these observations suggest that the regulation of global DNA methylation functions similarly in stable und unstable lines, regardless of the imprinting phenotype, and the cause of *Dlk1-Dio3* instability appears to be more targeted.

### Genetic background predetermines imprint stability in pluripotent cells

To account for the fundamental differences in imprint stability between pluripotent cell lines we considered two options – the stabilization of a random epigenetic event occurring in the embryo or, alternatively, predetermination by genetic factors. We turned our attention to the latter possibility, given that the *Dlk1* reporter alleles had been generated in F1 ESCs derived from a B6J x 129 cross and, following strain establishment, the mice were maintained on a mixed background. We derived ESC lines with pure B6J and 129 genetic backgrounds and investigated their imprinting status by bisulfite sequencing. This revealed that B6J ESCs had IG-DMR methylation levels strikingly similar to unstable reporter lines and consistent with LOI (Fig. 4A). In contrast, 129 lines displayed MOI methylation levels (Fig. 4A). The impact of genetic background on imprint stability was confirmed in context of the reporter alleles by crossing *Dlk1*^*Tomato*^ females with B6J and 129 males (Fig. S3A).

**Fig. 4.**
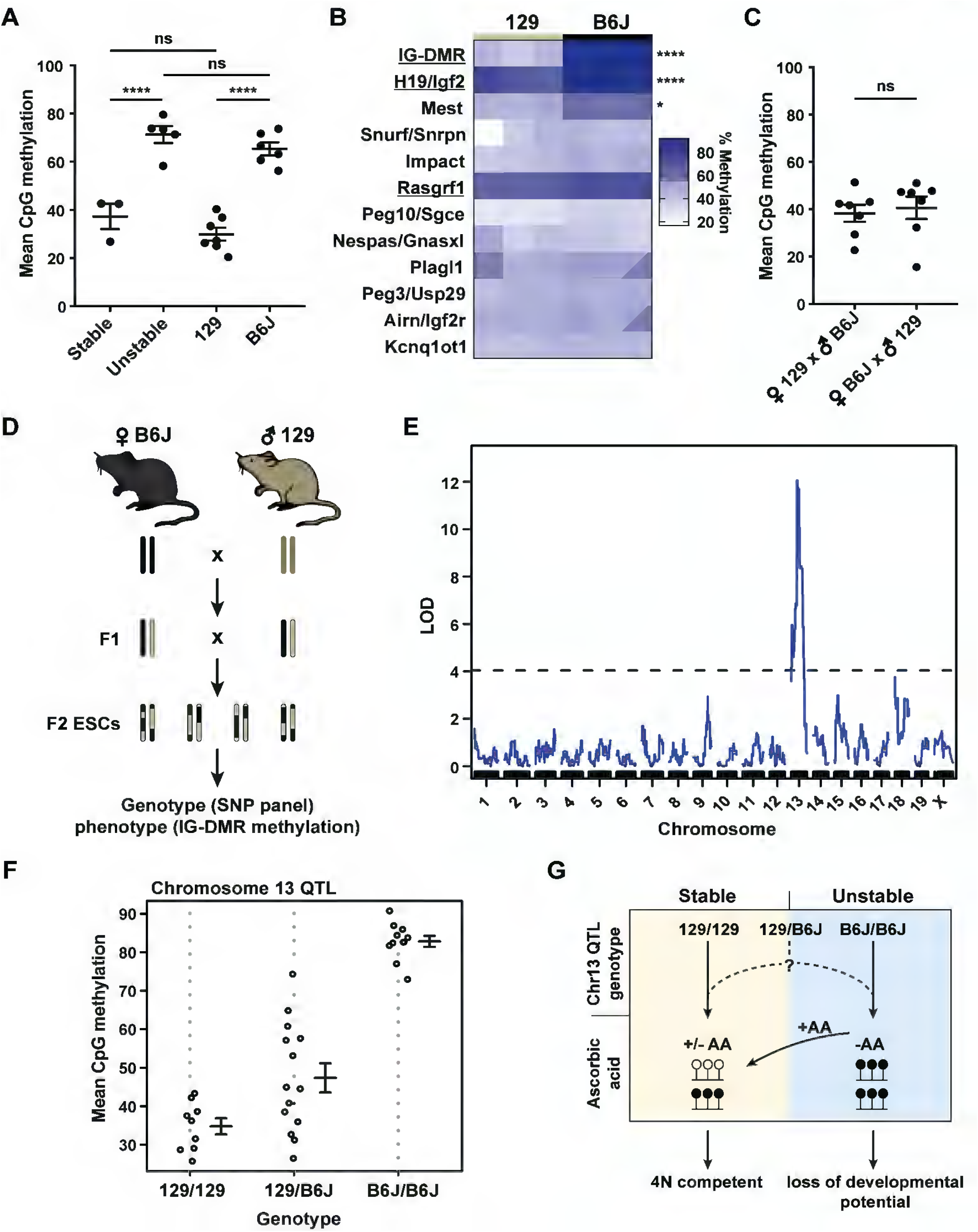
Genetic background predetermines imprint stability in pluripotent cells. (A) DNA methylation at the IG-DMR in stable and unstable reporter ESCs and ESCs with pure 129 and B6 backgrounds at p6. (B) DNA methylation at indicated germline ICRs in 129 and B6J ESCs. Each column represents an ESC line at p15. Paternal ICRs are underlined. ICRs are ranked by adjusted p-value and significance calculated with a two-way ANOVA with Sidak’s multiple comparison test. (****) indicates p<0.0001 and (*) p<0.05. (C) IG-DMR methylation in ESCs derived by reciprocal F1 crosses between 129 and B6J. Bars in (A,C) indicate mean and standard error. (****) indicates p<0.0001 and (ns) not significant with a one-way ANOVA and Turkey’s multiple comparison test. (D) Schematic of QTL analysis approach. (E) Single-QTL analysis of F2 lines. Y-axis shows the logarithm of the odds (LOD) score. Dashed line indicates the genome-wide LOD threshold cutoff of alpha = 0.1. (F) Main effect plot of the QTL max (R/qtl) shows the genotype at the chromosome 13 QTL (x-axis) for each F2 ESC line and the corresponding phenotype (y-axis; mean CpG methylation at the IG-DMR). 129/129 indicates homozygous for 129, 129/B6J heterozygous and B6J/B6J homozygous for B6J. Error bars represent mean with standard error. (G) Model for imprint stability in the context of chromosome 13 genetics: ESC lines with a homozygous 129 genotype (129/129) at the chromosome 13 QTL are stable, display MOI in the presence or absence of ascorbic acid (+/−AA) and are 4N competent. Lines with a homozygous B6J genotype at the QTL (B6J/B6J) are unstable and have LOI and loss of developmental potential in the absence of ascorbic acid (−AA) but LOI can be prevented by addition of ascorbic acid (+AA) to the cell culture media. Heterozygous lines (129/B6J) can adopt either stable or unstable trajectories, which is likely to be dependent on the inheritance of least one background-dependent secondary regulator (indicated by ‘?’).

To determine whether imprint instability was restricted to the IG-DMR, we analyzed a total of 12 germline ICRs in 129 and B6J ESCs by targeted next-generation bisulfite sequencing (tNGBS). While most ICRs exhibited similar DNA methylation levels in 129 and B6J ESCs, the *Igf2-H19* and *Mest* loci also had significantly increased ICR methylation levels in the B6J background (Fig. 4B). This supports the notion that the background-associated epigenetic instability we observed impacts specific susceptible loci, while the remainder of ICRs remain unaffected.

Of note, while neither B6J nor 129 mouse embryonic fibroblasts (MEFs) exhibited IG-DMR hypermethylation, derivative B6J iPSCs acquired LOI during reprogramming, whereas 129 iPSCs did not (Fig. S3B). Addition of AA during reprogramming prevented LOI in early passage B6J iPSCs (p1; Fig. S3C). However, B6J, but not 129, iPSCs acquired LOI upon continued passaging in absence of AA (Fig. S3C), mirroring our observations in ESCs. Together, these observations suggest that susceptibility of the B6J background to LOI at *Dlk1-Dio3* affects both established pluripotent cells and cells undergoing reprogramming, but does not arise in the developing embryo (Swanzey and Stadtfeld, 2016) or derivative MEFs.

### QTL analysis reveals a causal region on chromosome 13

To better understand the heredity of the imprinting phenotype, we analyzed ESC lines generated from B6Jx129 F1 crosses. This revealed moderately increased levels of IG-DMR methylation at higher passages (Fig. S3D), indicating incomplete dominance of the stable phenotype. Importantly, F1 ESCs derived from reciprocal crosses were not significantly different (Fig. 4C), suggesting that the cause of the instability was neither due to an imprinted gene nor to a mutation within the *Dlk1-Dio3* locus itself.

To narrow down the genomic region(s) of interest, we conducted Quantitative Trait Locus (QTL) analysis on 34 F2 ESC lines utilizing a SNP genotyping array of 143,000 markers (Morgan et al., 2015) (Fig. 4D). F2 lines displayed a range of IG-DMR phenotypes that resembled 129, B6J and F1 ESC lines (Fig. S3E and Table S1). QTL analysis mapped the IG-DMR methylation phenotype to a single interval of approximately 15 MB on chromosome 13 (Fig. 4E). All cell lines with the 129/129 genotype at this region displayed methylation levels consistent with MOI, while all lines with the B6J/B6J genotype had LOI (Fig. 4F). This strongly suggests that the chromosome 13 QTL (which we will refer to as *Dlk1-Dio3* QTL) contains a factor that is the predominant cause of the IG-DMR phenotype. In addition, the overall phenotype distribution was bimodal, with the IG-DMR methylation of the majority of the lines being at the extremes (Fig. S3F), which is consistent with the profile of a monogenic trait. However, cells that are 129/B6J at the *Dlk1-Dio3* QTL can have either MOI methylation or increased levels consistent with LOI (Fig. 4F), suggesting the existence of secondary regulator(s) outside of this QTL that determine the stability in heterozygotes. No significant secondary QTL was identified, which may suggest the presence of multiple secondary QTLs with individual logarithm of the odds (LOD) values below the significance threshold achievable with the limited number of lines. A model of our current understanding of how the genetic composition of chromosome 13, together with the availability of AA, determines imprint stability and developmental potential of pluripotent cells is depicted in Fig. 4G.

To address whether imprint stability in 129 or, alternatively, imprint instability in B6J is a unique feature of these strains, we derived ESCs from C57BL/6NJ (B6N) mice, which are closely related to B6J (Keane et al., 2011). Surprisingly, IG-DMR methylation levels in B6N ESCs were consistent with MOI (Fig. S3G,H). As B6J is the founder strain from which B6N was established it is likely that, following strain separation, B6J either lost a factor that protects against DNA hypermethylation at *Dlk1-Dio3* or gained a factor that drives hypermethylation. Alternatively, 129 and B6N would have had to both acquire protective mutations separately. The similarity of 129 and B6N phenotypes, including the F1s (Fig. S3D, H), suggests that neither strain carries the B6J-associated variant that causes imprint instability.

Given the similarity of B6N and B6J genetics (Simon et al., 2013), we attempted to assemble a list of distinguishing variants within the QTL. However, this effort was hampered by the prevalence of low-confidence base calls and large gaps in the B6N genome assembly (Fig. S4). As similar issues afflict the current 129 assembly, additional efforts to complete annotations of the 129 and B6N genome may be required before it will be possible to identify causal structural and/or regulatory variants within the *Dlk1-Dio3* QTL. Of note, inspection of the chromosomal territory covered by the QTL revealed no known regulator of imprinting. However, the QTL overlaps with a conserved cluster of more than twenty KRAB zinc-finger proteins some of which implicated in sex-specific gene expression (Krebs et al., 2005) and at least two genomic intervals that have previously been linked to disease and developmental phenotypes (Kano et al., 2007; Treger et al., 2019; Wang et al., 2002) (Fig. S4). This raises the possibility that the variant(s) modulating imprinting stability may also have regulatory functions in other biological contexts.

## Discussion

Our characterization of *Dlk1-Dio3* imprint stability strongly suggests that locus-specific detrimental epigenetic abnormalities can arise in pluripotent cells as a consequence of defined genetic variants. To our knowledge the results described here are the first proof-of-principle of a genetic determinant of LOI in pluripotent cells and highlight the importance of considering the interplay between environmental conditions and genetic variation in determining the epigenetic landscape of a cell. We anticipate that challenging current stochastic models for the occurrence of LOI in pluripotent cells will incentivize studies to systematically identify genetic factors controlling the epigenetic stability of ESCs and iPSCs. Pluripotent cells derived from well-characterized inbred mouse strains (Skelly et al., 2019) may serve as a promising avenue to aid in the pursuit of this goal, and large pools of genetically heterogenous human pluripotent cell lines should in principle also be amenable to genetic analysis.

Our findings have ramifications for several unexplained observations relating to stem cell pluripotency. For example, the observed epigenetic instability of B6J ESCs provides a molecular explanation for the reported difficulties in establishing ESCs with sustained developmental potential from C57BL/6 mice (Auerbach et al., 2000; Kumagai et al., 2017), even though the use of this nomenclature does not allow us to unambiguously determine whether B6J mice or a related strain were used in these studies. Similarly, the superior epigenetic stability of 129 iPSCs might explain discrepancies in the reported frequencies of *Dlk1-Dio3* LOI during cellular reprogramming of cells on undefined backgrounds (Carey et al., 2011) and the high success rate of 4N complementation with iPSCs derived in basal reprogramming conditions in some studies (Boland et al., 2009). This stresses the need to consider genetic background as a strong modulator of fundamental stem cell properties.

Similar to our observations with murine ESCs, the *H19/IGF2* and *DLK1-DIO3* loci have also been shown to be particular prone to imprint loss in human ESCs (Kim et al., 2007; Rugg-Gunn et al., 2007). This raises the possibility that at least some of the factors or pathways controlling epigenetic stability at these two paternally-imprinted gene loci are conserved between mouse and human. While limitations in the current mouse genome assemblies prevented us from identifying the causal variant(s) encoded by the *Dlk1-Dio3* QTL, we speculate that the relevant genes may facilitate the recruitment of the *de novo* DNA methylation machinery to susceptible gene loci. This appears to be a dynamic process, potentially resembling recent observations made at pluripotency-associated enhancers (Song et al., 2019), that can be counteracted by stimulating DNA and/or histone demethylases. The ability to suppress imprint instability by ascorbic acid may prove beneficial when attempting to preserve the developmental potential of pluripotent cell lines that have been determined to be susceptible to ICR hypermethylation due to their genetic composition. In summary, our findings stress the importance of genetic determinants in regulating epigenetic stability and provide a strong rationale to further dissect the interplay between genetic variants, different culture conditions and specific cell states in controlling the epigenome. This may yield novel methods to reliably identify and maintain pluripotent cell lines with desirable epigenetic properties.

## Acknowledgements

We would like to thank the Sfeir lab, particularly Marco Tigano, Aleks Penev and Agnel Sfeir, for advice, discussion and sharing of reagents. We would also like to thank Sang Yong Kim for blastocyst injections, Paul Zumbo for help with bioinformatic analysis and Boaz Eliezer Aronson for providing virus for reprogramming experiments. We are thankful to Konrad Hochedlinger for comments on the manuscript and to all members of the Stadtfeld and Apostolou labs for comments and discussion.

## Author Contributions

E.S. and M.S. conceived and designed experiments. E.S. conducted and analyzed most experiments. M.T., T.F.M. and E.S. performed and analyzed TLC experiment. M.S. performed iPSC reprogramming experiments. E.A. provided reagents and helped with data acquisition. E.A. and M.S. acquired funding. E.S. and M.S. wrote the manuscript with input from E.A., M.T. and T.F.M.

## Declaration of Interests

Nothing to declare.

## Methods

### Mice

Experiments were conducted by deriving ESC lines from the following strains: C57BL/6NJ (Jackson 005304), C57BL/6J (Jackson 000664), 129S2 (Charles River 476), B6129SF1/J (Jackson 101043; a F1 cross between C57BL/6J and 129S1/SvImJ), *Dlk1*^*Tomato*^ and *Dlk1*^*Venus*^ (Swanzey et al. 2016). B6D2F1/J (Jackson 100006) females were used for 4N blastocyst injection experiments (see below). All animal experiments were done in accordance with the guidelines of the NYU School of Medicine IACUC.

### 4N Blastocyst Complementation

Zygotes were isolated from B6D2F1/J females and cultured overnight until they reached the 2-cell stage, as previously described. Embryos were electro-fused and those with successful fusion were cultured for another 2 days, followed by injection with indicated male ESC lines (Amlani et al., 2018). Viable pups were defined as those that survived to the indicated mid-gestation time point (E14.5 or E15.5) and were quantified as a percentage of the total number of blastocysts transferred into the recipient female (20 each).

### ESC Derivation and Culture

Each ESC line was derived from an individual E3.5 blastocyst as previously described (Czechanski et al., 2014). In brief, blastocysts were isolated and cultured in KnockOut DMEM (Gibco) supplemented with 15% KnockOut Serum Replacement (Gibco), 1% FBS, L-Glutamine, sodium pyruvate, penicillin, streptomycin, non-essential amino acids, 2-Mercaptoethanol, 1000 U/ml LIF, 1 μM PD03259010 and 3 μM CHIR99021 on a layer of Mitomycin-C-treated feeders for four days. Media was then changed to KO-DMEM with 15% FBS, L-Glutamine, penicillin-streptomycin, non-essential amino acids, 2-Mercaptoethanol and 1000 U/ml LIF (ESC medium) on Mitomycin-C-treated feeder cells. If applicable, L-Ascorbic acid (50 μg/ml) was added. For generation of clonal lines, ESCs were single cell sorted in 96-well plates using a MoFlo XDP cell sorter (Beckman Coulter) and subsequently expanded.

### MEF Derivation and iPSC Reprogramming

MEF cultures were established by trypsin digestion of midgestation (E14.5–E15.5) 129S2 and C57BL/6J embryos and maintained in DMEM supplemented with 10% FBS, L-glutamine, penicillin/streptomycin, nonessential amino acids, and β-mercaptoethanol (MEF medium). Reprogramming was carried out by transducing MEFs with a doxycycline-inducible mouse OKSM (pHAGE2-tetO-STEMCCA)(Sommer et al., 2009) and rtTA (FUdeltaGW-rtTA, Addgene 19780)(Maherali et al., 2008) lentiviruses. Transduced MEFs were then seeded at a density of approximately 1,500 cells/cm^2^ onto a layer of Mitomycin-C-treated feeder cells in ESC media supplemented with 1 μg/mL doxycycline and, if applicable, L-ascorbic acid (50 μg/mL). Media was changed every two days and on day 11, doxycycline and ascorbic acid were withdrawn.

### Cell Differentiation and Flow Cytometry

Trypsinized ESCs were pre-plated for 30 minutes to remove feeder cells and seeded onto gelatinized plates at a density of 20,000 cells/cm^2^. The next day, the media was changed to MEF medium supplemented with 0.4 μg/ml retinoic acid. Media was changed daily for five days and imaged using a Nikon Eclipse TiE inverted microscope with filters to detect Venus (Ex 500/20; Em 535/30) and Tomato (Ex 545/30; Em 620/60). For quantification of *Dlk1* reporter expression, dissociated cultures were acquired on an LSRII cytometer (BD Biosciences) and analyzed with FlowJo software (Tree Star Inc.).

### Tissue Isolation and Reporter Detection

Mouse embryos isolated at E15.5 were imaged using a Nikon SMZ1500 Stereo Fluorescence Microscope. For flow cytometry, tissues were isolated in the same manner as MEF derivations (described above), followed by analysis on an LSRII cytometer (BD Biosciences) and analyzed with FlowJo software (Tree Star Inc.).

### DNA Methylation Analysis

Genomic DNA was isolated using proteinase K in lysis buffer, pH 8 (100 mM Tris-HCl, 5 mM EDTA, 0.2% SDS, 200 mM NaCl) and reconstituted in TE buffer, pH 7.5 (10 mM Tris-HCl, 1 mM EDTA). Bisulfite sequencing of the IG-DMR was conducted using the ADS1452-FS1/FS2/FS3 EpigenDx assays (chr12:109,528,253-109,528,471 in mm10). If indicated, a portion of this region was analyzed (ADS1452-FS3; chr12:109,528,391-109,528,471). Targeted Next Generation Bisulfite Sequencing was conducted by EpigenDx for the following germline ICR regions in mm10: *Nespas/Gnasxl* (chr2:174292903-174299786), *Peg10/Sgce* (chr6:4746303-4749370), *Mest* (chr6:30735840-30739965), *Peg3/Usp29* (chr7:6727356-6733209), *Snurf/Snrpn* (chr7:60003140-60005283), *Airn/Igf2r* (chr17:12741760-12742949), *Impact* (chr18:12972847-12974748), *H19/Igf2* (chr7:142579886-142582026), *Rasgrf1* (chr9:89869688-89878981), IG-DMR (chr12:109525755-109530399). Thin-layer chromatography (TLC) was conducted as previously described (Tahiliani et al., 2009).

### QTL Analysis

34 ESC lines from an C57BL/6J × 129S1/SvImJ F2 intercross were sequenced by Neogen with the Giga Mouse Universal Genotyping Array (GigaMUGA), which includes >143,000 total SNPs and >34,000 informative SNPs for C57BL/6J and 129S1/SvImJ strains. For the phenotype parameter, IG-DMR methylation was assessed in all lines by bisulfite sequencing. Single-locus QTL analysis was performed using R/qtl (Broman et al., 2003) with standard interval mapping with Haley Knott regression. The null distribution for the genome-wide LOD score was generated by performing 10,000 permutation tests. The p value for LOD peaks was then calculated using an alpha of 0.1. The location of the QTL interval was estimated using the Bayes 95% credible interval. The QTL data can be accessed in HaploQA (https://haploqa.jax.org/tag/f2.html).

## References

Amlani, B., Liu, Y., Chen, T., Ee, L.S., Lopez, P., Heguy, A., Apostolou, E., Kim, S.Y., and Stadtfeld, M. (2018). Nascent Induced Pluripotent Stem Cells Efficiently Generate Entirely iPSC-Derived Mice while Expressing Differentiation-Associated Genes. Cell Rep 22, 876–884.

Auerbach, W., Dunmore, J.H., Fairchild-Huntress, V., Fang, Q., Auerbach, A.B., Huszar, D., and Joyner, A.L. (2000). Establishment and chimera analysis of 129/SvEv- and C57BL/6-derived mouse embryonic stem cell lines. Biotechniques 29, 1024–1028, 1030, 1032.

Bar, S., and Benvenisty, N. (2019). Epigenetic aberrations in human pluripotent stem cells. EMBO J 38.

Blaschke, K., Ebata, K.T., Karimi, M.M., Zepeda-Martinez, J.A., Goyal, P., Mahapatra, S., Tam, A., Laird, D.J., Hirst, M., Rao, A., et al. (2013). Vitamin C induces Tet-dependent DNA demethylation and a blastocyst-like state in ES cells. Nature 500, 222–226.

Boland, M.J., Hazen, J.L., Nazor, K.L., Rodriguez, A.R., Gifford, W., Martin, G., Kupriyanov, S., and Baldwin, K.K. (2009). Adult mice generated from induced pluripotent stem cells. Nature 461, 91–94.

Broman, K.W., Wu, H., Sen, S., and Churchill, G.A. (2003). R/qtl: QTL mapping in experimental crosses. Bioinformatics 19, 889–890.

Cantone, I., and Fisher, A.G. (2013). Epigenetic programming and reprogramming during development. Nat Struct Mol Biol 20, 282–289.

Carey, B.W., Markoulaki, S., Hanna, J.H., Faddah, D.A., Buganim, Y., Kim, J., Ganz, K., Steine, E.J., Cassady, J.P., Creyghton, M.P., et al. (2011). Reprogramming factor stoichiometry influences the epigenetic state and biological properties of induced pluripotent stem cells. Cell Stem Cell 9, 588–598.

Choi, J., Huebner, A.J., Clement, K., Walsh, R.M., Savol, A., Lin, K., Gu, H., Di Stefano, B., Brumbaugh, J., Kim, S.Y., et al. (2017). Prolonged Mek1/2 suppression impairs the developmental potential of embryonic stem cells. Nature 548, 219–223.

Cimmino, L., Neel, B.G., and Aifantis, I. (2018). Vitamin C in Stem Cell Reprogramming and Cancer. Trends Cell Biol 28, 698–708.

Czechanski, A., Byers, C., Greenstein, I., Schrode, N., Donahue, L.R., Hadjantonakis, A.K., and Reinholdt, L.G. (2014). Derivation and characterization of mouse embryonic stem cells from permissive and nonpermissive strains. Nat Protoc 9, 559–574.

Dean, W., Bowden, L., Aitchison, A., Klose, J., Moore, T., Meneses, J.J., Reik, W., and Feil, R. (1998). Altered imprinted gene methylation and expression in completely ES cell-derived mouse fetuses: association with aberrant phenotypes. Development 125, 2273–2282.

Dirks, R.A.M., van Mierlo, G., Kerstens, H.H.D., Bernardo, A.S., Kobolak, J., Bock, I., Maruotti, J., Pedersen, R.A., Dinnyes, A., Huynen, M.A., et al. (2019). Allele-specific RNA-seq expression profiling of imprinted genes in mouse isogenic pluripotent states. Epigenetics Chromatin 12, 14.

Duffie, R., and Bourc’his, D. (2013). Parental epigenetic asymmetry in mammals. Curr Top Dev Biol 104, 293–328.

Humpherys, D., Eggan, K., Akutsu, H., Hochedlinger, K., Rideout, W.M., 3rd, Biniszkiewicz, D., Yanagimachi, R., and Jaenisch, R. (2001). Epigenetic instability in ES cells and cloned mice. Science 293, 95–97.

Johannesson, B., Sagi, I., Gore, A., Paull, D., Yamada, M., Golan-Lev, T., Li, Z., LeDuc, C., Shen, Y., Stern, S., et al. (2014). Comparable frequencies of coding mutations and loss of imprinting in human pluripotent cells derived by nuclear transfer and defined factors. Cell Stem Cell 15, 634–642.

Kano, H., Kurahashi, H., and Toda, T. (2007). Genetically regulated epigenetic transcriptional activation of retrotransposon insertion confers mouse dactylaplasia phenotype. Proc Natl Acad Sci U S A 104, 19034–19039.

Keane, T.M., Goodstadt, L., Danecek, P., White, M.A., Wong, K., Yalcin, B., Heger, A., Agam, A., Slater, G., Goodson, M., et al. (2011). Mouse genomic variation and its effect on phenotypes and gene regulation. Nature 477, 289–294.

Kim, K.P., Thurston, A., Mummery, C., Ward-van Oostwaard, D., Priddle, H., Allegrucci, C., Denning, C., and Young, L. (2007). Gene-specific vulnerability to imprinting variability in human embryonic stem cell lines. Genome Res 17, 1731–1742.

Krebs, C.J., Larkins, L.K., Khan, S.M., and Robins, D.M. (2005). Expansion and diversification of KRAB zinc-finger genes within a cluster including Regulator of sex-limitation 1 and 2. Genomics 85, 752–761.

Kumagai, K., Takanashi, M., Ohno, S.I., Kuroda, M., and Sudo, K. (2017). An improved Red/ET recombineering system and mouse ES cells culture conditions for the generation of targeted mutant mice. Exp Anim 66, 125–136.

Liu, L., Luo, G.Z., Yang, W., Zhao, X., Zheng, Q., Lv, Z., Li, W., Wu, H.J., Wang, L., Wang, X.J., et al. (2010). Activation of the imprinted Dlk1-Dio3 region correlates with pluripotency levels of mouse stem cells. J Biol Chem 285, 19483–19490.

Maherali, N., Ahfeldt, T., Rigamonti, A., Utikal, J., Cowan, C., and Hochedlinger, K. (2008). A high-efficiency system for the generation and study of human induced pluripotent stem cells. Cell Stem Cell 3, 340–345.

Mo, C.F., Wu, F.C., Tai, K.Y., Chang, W.C., Chang, K.W., Kuo, H.C., Ho, H.N., Chen, H.F., and Lin, S.P. (2015). Loss of non-coding RNA expression from the DLK1-DIO3 imprinted locus correlates with reduced neural differentiation potential in human embryonic stem cell lines. Stem Cell Res Ther 6, 1.

Monfort, A., and Wutz, A. (2013). Breathing-in epigenetic change with vitamin C. EMBO Rep 14, 337–346.

Morgan, A.P., Fu, C.P., Kao, C.Y., Welsh, C.E., Didion, J.P., Yadgary, L., Hyacinth, L., Ferris, M.T., Bell, T.A., Miller, D.R., et al. (2015). The Mouse Universal Genotyping Array: From Substrains to Subspecies. G3 (Bethesda) 6, 263–279.

Pastor, W.A., Chen, D., Liu, W., Kim, R., Sahakyan, A., Lukianchikov, A., Plath, K., Jacobsen, S.E., and Clark, A.T. (2016). Naive Human Pluripotent Cells Feature a Methylation Landscape Devoid of Blastocyst or Germline Memory. Cell Stem Cell 18, 323–329.

Rugg-Gunn, P.J., Ferguson-Smith, A.C., and Pedersen, R.A. (2007). Status of genomic imprinting in human embryonic stem cells as revealed by a large cohort of independently derived and maintained lines. Hum Mol Genet 16 Spec No. 2, R243–251.

Sanli, I., and Feil, R. (2015). Chromatin mechanisms in the developmental control of imprinted gene expression. Int J Biochem Cell Biol 67, 139–147.

Simon, M.M., Greenaway, S., White, J.K., Fuchs, H., Gailus-Durner, V., Wells, S., Sorg, T., Wong, K., Bedu, E., Cartwright, E.J., et al. (2013). A comparative phenotypic and genomic analysis of C57BL/6J and C57BL/6N mouse strains. Genome Biol 14, R82.

Skelly, D.A., Czechanski, A., Byers, C., Aydin, S., Spruce, C., Olivier, C., Choi, K., Gatti, D.M., Raghupathy, N.M., Stanton, A., et al. (2019). Genetic variation influences pluripotent ground state stability in mouse embryonic stem cells through a hierarchy of molecular phenotypes. bioRxiv, 552059.

Sommer, C.A., Stadtfeld, M., Murphy, G.J., Hochedlinger, K., Kotton, D.N., and Mostoslavsky, G. (2009). Induced pluripotent stem cell generation using a single lentiviral stem cell cassette. Stem Cells 27, 543–549.

Song, Y., van den Berg, P.R., Markoulaki, S., Soldner, F., Dall’Agnese, A., Henninger, J.E., Drotar, J., Rosenau, N., Cohen, M.A., Young, R.A., et al. (2019). Dynamic Enhancer DNA Methylation as Basis for Transcriptional and Cellular Heterogeneity of ESCs. Mol Cell 75, 905–920 e906.

Stadtfeld, M., Apostolou, E., Akutsu, H., Fukuda, A., Follett, P., Natesan, S., Kono, T., Shioda, T., and Hochedlinger, K. (2010). Aberrant silencing of imprinted genes on chromosome 12qF1 in mouse induced pluripotent stem cells. Nature 465, 175–181.

Stadtfeld, M., Apostolou, E., Ferrari, F., Choi, J., Walsh, R.M., Chen, T., Ooi, S.S., Kim, S.Y., Bestor, T.H., Shioda, T., et al. (2012). Ascorbic acid prevents loss of Dlk1-Dio3 imprinting and facilitates generation of all-iPS cell mice from terminally differentiated B cells. Nat Genet 44, 398–405, s391-392.

Stelzer, Y., Shivalila, C.S., Soldner, F., Markoulaki, S., and Jaenisch, R. (2015). Tracing dynamic changes of DNA methylation at single-cell resolution. Cell 163, 218–229.

Swanzey, E., and Stadtfeld, M. (2016). A reporter model to visualize imprinting stability at the Dlk1 locus during mouse development and in pluripotent cells. Development 143, 4161–4166.

Tahiliani, M., Koh, K.P., Shen, Y., Pastor, W.A., Bandukwala, H., Brudno, Y., Agarwal, S., Iyer, L.M., Liu, D.R., Aravind, L., et al. (2009). Conversion of 5-methylcytosine to 5-hydroxymethylcytosine in mammalian DNA by MLL partner TET1. Science 324, 930–935.

Tee, W.W., and Reinberg, D. (2014). Chromatin features and the epigenetic regulation of pluripotency states in ESCs. Development 141, 2376–2390.

Treger, R.S., Pope, S.D., Kong, Y., Tokuyama, M., Taura, M., and Iwasaki, A. (2019). The Lupus Susceptibility Locus Sgp3 Encodes the Suppressor of Endogenous Retrovirus Expression SNERV. Immunity 50, 334–347 e339.

Wang, D., Lemon, W.J., and You, M. (2002). Linkage disequilibrium mapping of novel lung tumor susceptibility quantitative trait loci in mice. Oncogene 21, 6858–6865.

Yagi, M., Kishigami, S., Tanaka, A., Semi, K., Mizutani, E., Wakayama, S., Wakayama, T., Yamamoto, T., and Yamada, Y. (2017). Derivation of ground-state female ES cells maintaining gamete-derived DNA methylation. Nature 548, 224–227.

