## Supplemental Figures 1-4 and Table S1 for "A susceptibility locus on chromosome 13 profoundly impacts the stability of genomic imprinting in mouse pluripotent stem cells"

Swanzey_FigS1

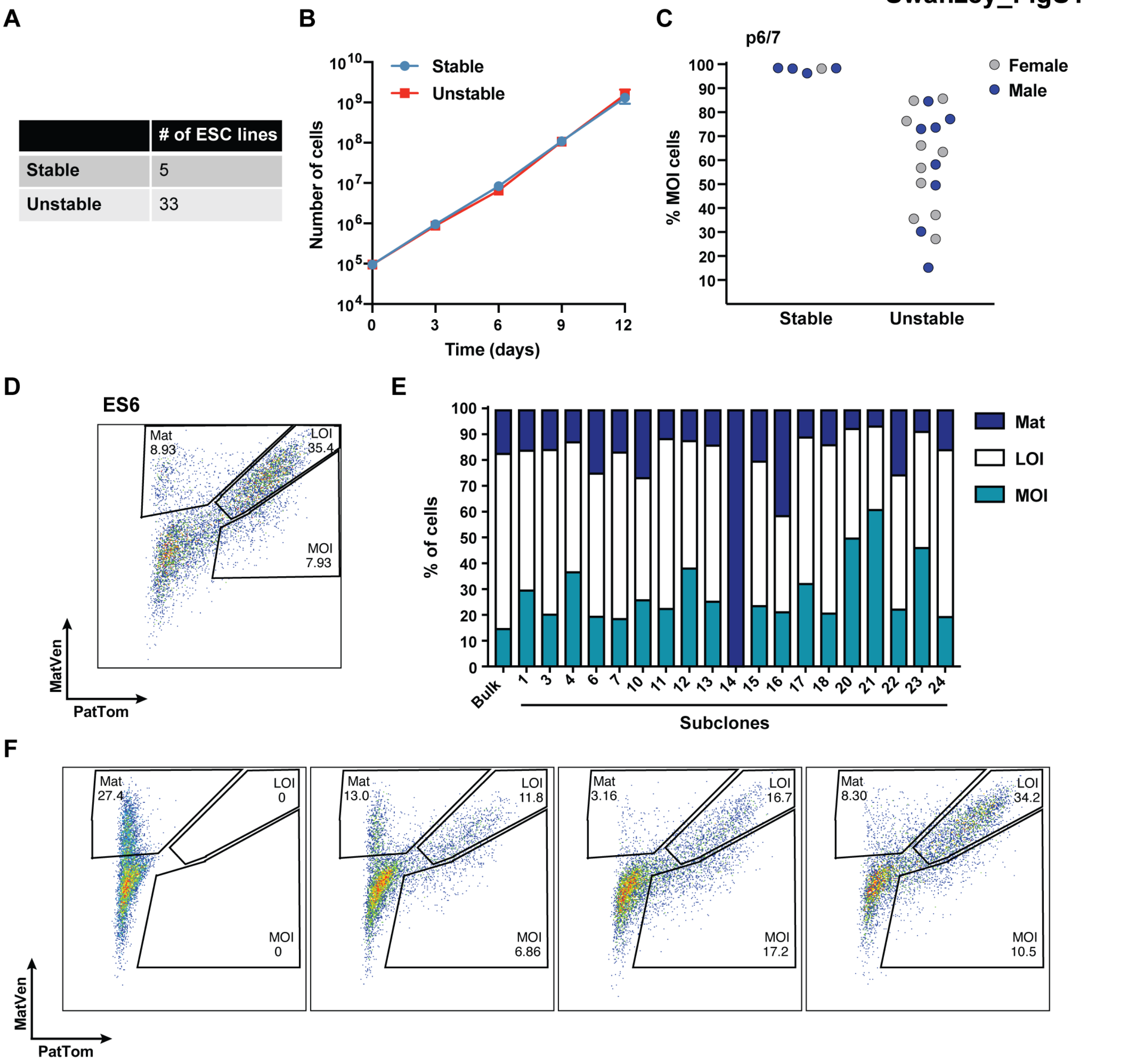


**Fig. S1. Characterization of stable and unstable reporter ESC lines (related to Figure 2).** (A) In total, 5 stable and 33 unstable reporter lines were identified and characterized based on whether MOI decreases ("unstable") or remains high ("stable") during continued passaging. (B) Growth curves for representative stable and unstable reporter lines over 12 days, with passaging every three days. Error bars represent standard deviation of 3 biological replicates, with each being the mean value of 3 technical replicates. (C) Quantification of the percentage of cells with MOI in subset of lines at p6 or p7. Female lines are indicated in gray, male lines are in blue. (D) Flow cytometry profile of ES6, a *Dlk1*$^{PatTom/MatVen}$ ESC line that shows high LOI in addition to maternal-only (Mat) expression at p8. (E) ES6 was single-cell sorted into 96-well plates at p4 and subclones were expanded four more passages for analysis by flow cytometry. Bar graphs show the quantification of the percentage of *Dlk1*-expressing cells that have LOI, MOI or Mat reporter fluorescence, compared to the bulk parental line. (F) Representative flow cytometry profiles of ES6 subclones with a range of phenotypes.

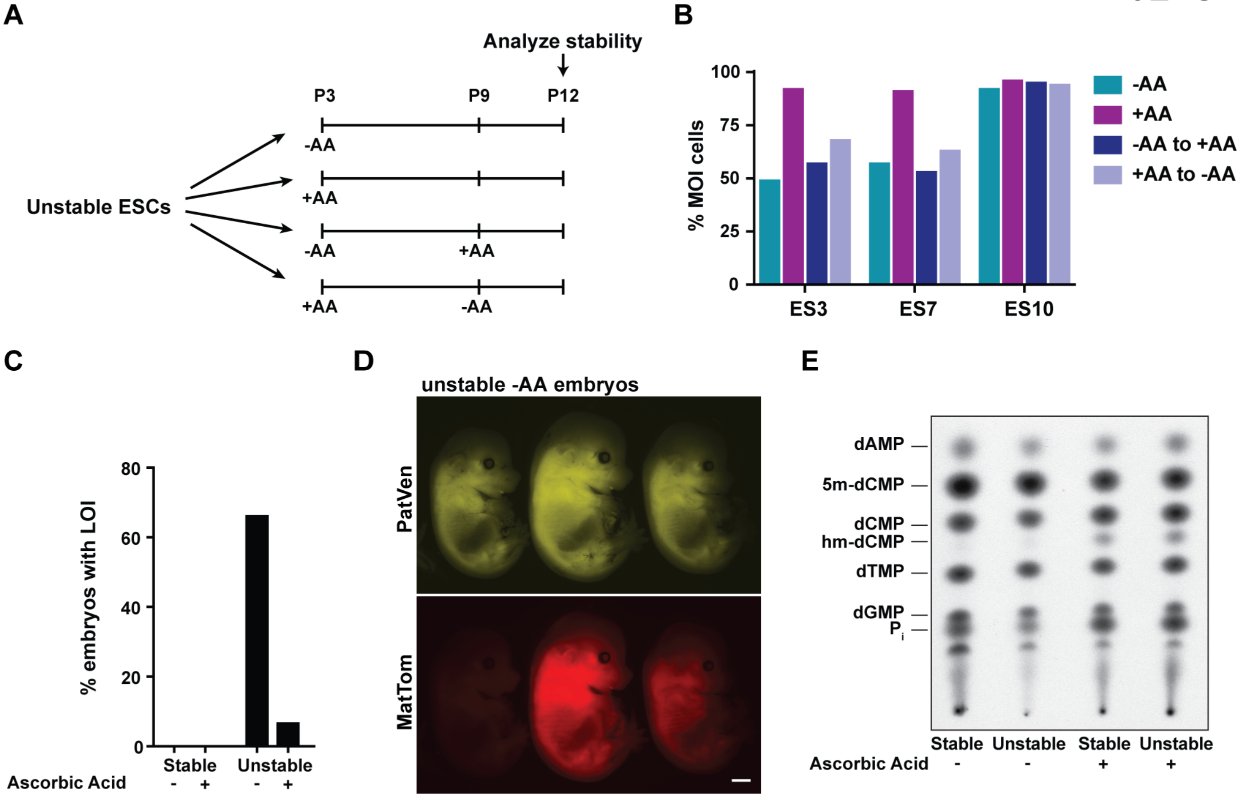


**Fig. S2. AA treatment of unstable lines attenuates imprinting instability (related to Figure 3).** (A) Experimental schematic. Unstable ESC lines were split at early passage, to be continued in the same conditions (-AA), treated with AA (+AA) continually or for the AA treatment to be changed at P9. Imprint stability was then assessed at p12. (B) Quantification of the percentage of *Dlk1*-expressing cells with MOI fluorescence in two unstable (ES3 and ES7) lines and one stable (ES10) line in the indicated conditions. (C) Percentage of viable embryos from 4N injections that displayed fluorescence indicative of *Dlk1* LOI. Unstable ESCs (-AA): 2/3 embryos had LOI. Unstable ESCs (+AA): 1/14 embryos had LOI. Stable ESCs (-AA): 0/11 embryos had LOI. Stable ESCs (+AA): 0/16 had LOI. (D) Fluorescent images of *Dlk1*$^{PatVen/MatTom}$ embryos obtained after 4N injections with unstable ESCs cultured in absence of AA. ESCs were injected at p7 and the embryos were dissected at E14.5. From left to right: embryo displaying MOI, embryo with LOI, embryo with LOI. Scale bar = 1mm. (E) TLC blot to assess global methylation of stable and unstable lines that were cultured in the presence or absence of AA from p2 until the samples were taken for analysis (p7).

Swanzey_FigS3

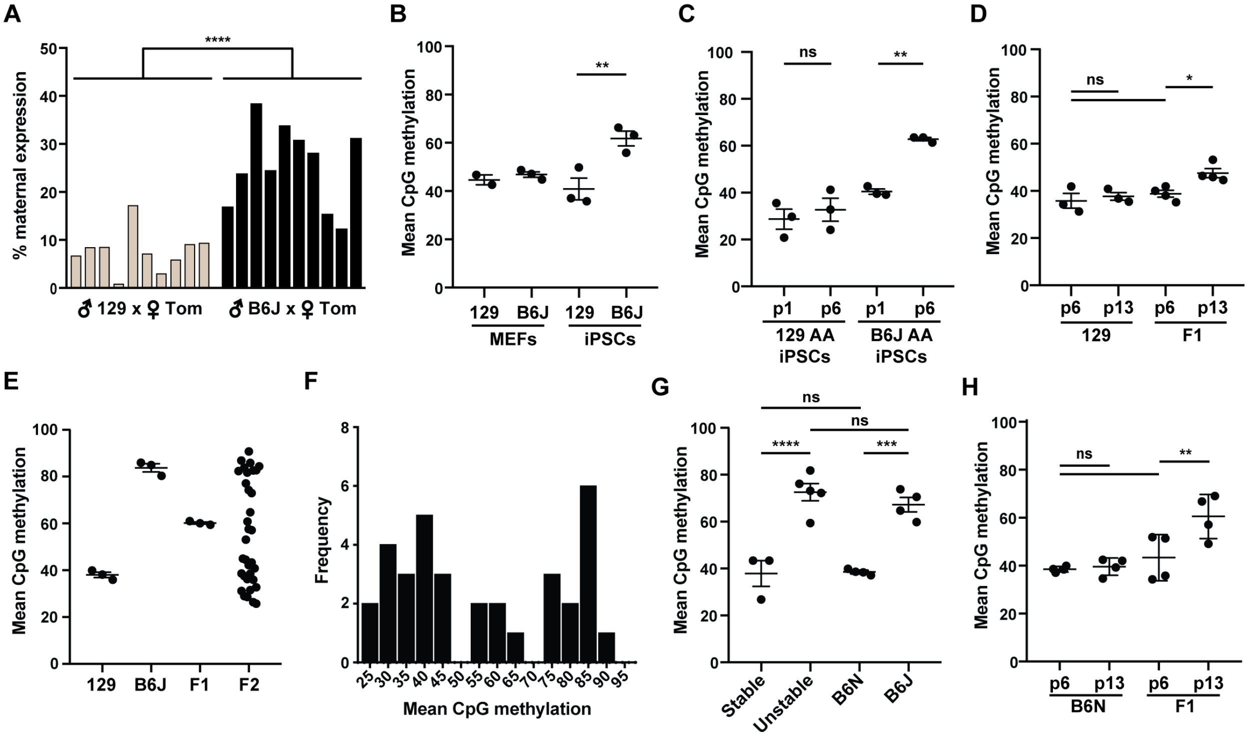


**Fig. S3. Genetic background determines imprint stability at *Dlk1-Dio3* (related to Figure 4).** (A) Percentage of F1 ESCs derived by crossing B6J or 129 males with $Dlk1^{Tomato}$ females that expressed the maternally-inherited reporter allele. Each bar represents one ESC line. (****) indicates p<0.0001 with a one-way ANOVA and Turkey's multiple comparison test. (B) IG-DMR methylation (EpigenDx assay ADS1452-FS3) in 129 and B6J MEFs and p1 iPSCs derived in absence of AA. (C) IG-DMR methylation in 129 and B6J iPSCs reprogrammed in the presence of AA and then cultured without AA until p1 and p6. (**) indicates p<0.01, (ns) not significant, with a one-way ANOVA and Turkey's multiple comparison test. (D) IG-DMR methylation in 129 and F1 (129 x B6J) ESC lines at p6 and p13. (E) IG-DMR methylation in 129, B6J, F1 and 34 F2 ESCs at p15. Error bars represent mean with standard error. (F) IG-DMR phenotype distribution frequency of F2 samples. (G) Mean CpG methylation at the IG-DMR (EpigenDx assay ADS1452-FS3) in stable and unstable reporter ESC lines and lines with B6N or B6J backgrounds at p6. (H) IG-DMR methylation in pure B6N and F1 (B6N x B6J) lines at p6 and p13. Bars represent mean and standard error. (****) indicates p<0.0001, (***) p<0.001, (**) p<0.01, (ns) not significant, with a one-way ANOVA and Turkey's multiple comparison test.

**Swanzey_FigS4**

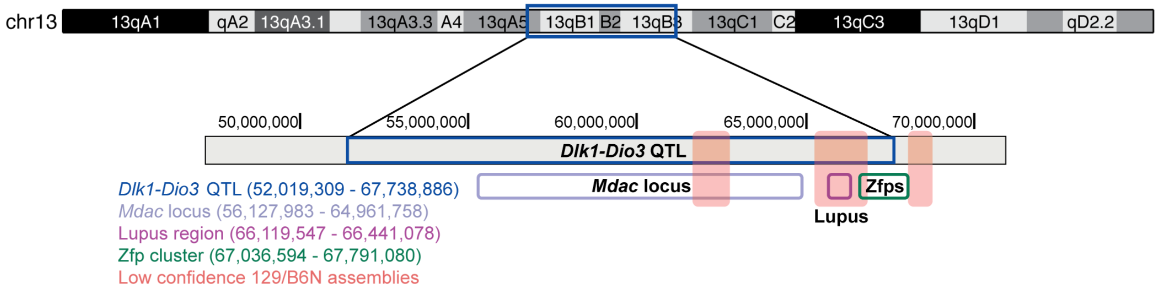


**Fig. S4. Features of the *Dlk1-Dio3* QTL on chromosome 13 (related to Figure 4).** Schematic of chromosome 13 (chr13) with indicated cytogenic bands. The QTL interval, estimated using the Bayes 95% credible interval, is outlined in blue (top). Higher resolution schematic of the region around the *Dlk1-Dio3* QTL with select overlapping genomic features and their positions (mm10) highlighted (bottom). Chosen features include the genomic region to which the unidentified *modifier of Dac* (*Mdac*) gene has been mapped, a region with gene loci conferring susceptibility to lupus and an area encoding 25 KRAB zinc-finger proteins (ZFPs). Red-shaded bars indicated areas where the current 129 and B6N genome annotations are less reliable.

**Table S1. F2 ESCs for QTL analysis** (related to Figure 4)

| ID# | ESC_Line_Name | Cross | Sex | Passage | IG-DMR_mean_methylation |
|---|---|---|---|---|---|
| 163 | 032919_F2_ES1 | B6129SF1/J x B6129SF1/J | F | 15 | 29.0 |
| 164 | 032919_F2_ES2 | B6129SF1/J x B6129SF1/J | M | 15 | 74.3 |
| 165 | 032919_F2_ES3 | B6129SF1/J x B6129SF1/J | M | 15 | 53.1 |
| 166 | 032919_F2_ES4 | B6129SF1/J x B6129SF1/J | F | 15 | 90.7 |
| 167 | 032919_F2_ES5 | B6129SF1/J x B6129SF1/J | F | 15 | 42.2 |
| 168 | 032919_F2_ES6 | B6129SF1/J x B6129SF1/J | F | 15 | 84.4 |
| 169 | 032919_F2_ES7 | B6129SF1/J x B6129SF1/J | F | 15 | 32.7 |
| 170 | 032919_F2_ES8 | B6129SF1/J x B6129SF1/J | M | 15 | 43.3 |
| 171 | 032919_F2_ES9 | B6129SF1/J x B6129SF1/J | F | 15 | 26.4 |
| 172 | 032919_F2_ES10 | B6129SF1/J x B6129SF1/J | F | 15 | 31.5 |
| 173 | 032919_F2_ES11 | B6129SF1/J x B6129SF1/J | F | 15 | 60.8 |
| 174 | 032919_F2_ES12 | B6129SF1/J x B6129SF1/J | M | 15 | 83.8 |
| 175 | 032919_F2_ES13 | B6129SF1/J x B6129SF1/J | M | 15 | 86.9 |
| 176 | 032919_F2_ES14 | B6129SF1/J x B6129SF1/J | M | 15 | 57.1 |
| 177 | 032919_F2_ES15 | B6129SF1/J x B6129SF1/J | M | 15 | 82.6 |
| 178 | 032919_F2_ES16 | B6129SF1/J x B6129SF1/J | F | 15 | 31.2 |
| 179 | 032919_F2_ES17 | B6129SF1/J x B6129SF1/J | M | 15 | 37.5 |
| 180 | 032919_F2_ES18 | B6129SF1/J x B6129SF1/J | M | 15 | 81.7 |
| 181 | 032919_F2_ES19 | B6129SF1/J x B6129SF1/J | F | 15 | 25.7 |
| 182 | 032919_F2_ES20 | B6129SF1/J x B6129SF1/J | F | 15 | 77.1 |
| 183 | 032919_F2_ES21 | B6129SF1/J x B6129SF1/J | M | 15 | 44.5 |
| 184 | 032919_F2_ES22 | B6129SF1/J x B6129SF1/J | F | 15 | 36.0 |
| 185 | 032919_F2_ES23 | B6129SF1/J x B6129SF1/J | F | 15 | 45.0 |
| 186 | 032919_F2_ES24 | B6129SF1/J x B6129SF1/J | M | 15 | 64.8 |
| 187 | 032919_F2_ES25 | B6129SF1/J x B6129SF1/J | F | 15 | 40.8 |
| 188 | 032919_F2_ES26 | B6129SF1/J x B6129SF1/J | M | 15 | 57.6 |
| 189 | 032919_F2_ES27 | B6129SF1/J x B6129SF1/J | F | 15 | 38.6 |
| 190 | 032919_F2_ES28 | B6129SF1/J x B6129SF1/J | M | 15 | 82.7 |
| 191 | 032919_F2_ES29 | B6129SF1/J x B6129SF1/J | M | 15 | 36.2 |
| 192 | 032919_F2_ES30 | B6129SF1/J x B6129SF1/J | F | 15 | 28.6 |
| 193 | 032919_F2_ES31 | B6129SF1/J x B6129SF1/J | M | 15 | 38.5 |
| 194 | 032919_F2_ES32 | B6129SF1/J x B6129SF1/J | F | 15 | 85.9 |
| 195 | 032919_F2_ES33 | B6129SF1/J x B6129SF1/J | M | 15 | 82.4 |
| 196 | 032919_F2_ES34 | B6129SF1/J x B6129SF1/J | F | 15 | 73.0 |
